# Comparative assessment of computed tomography and magnetic resonance imaging of the spider morph of *Python regius* and wild type *Python regius* to evaluate the morphological correlate of the wobble syndrome

**DOI:** 10.1101/2022.03.24.485672

**Authors:** Fabian Schrenk, Matthias Starck, Ingmar Kiefer, Thomas Flegel, Wiebke Tebrün, Michael Pees

## Abstract

There is general awareness of artificial selection and its potential implications on health and welfare of animals. Despite growing popularity and increasing numbers of breeds of atypical colour and pattern variants in reptiles, only few studies have investigated the appearance and cause of various diseases associated with colour morphs. Ball pythons (*Python regius*) are among the most frequently bred reptiles and breeders selected for a multitude of different colour and pattern morphs. Among those colour variants, the spider morph of the ball python is frequently associated with the wobble syndrome. The aim of this study was to determine, whether a morphological variant can be found and brought in association with the clinical occurrence of the wobble syndrome in spider ball pythons, using MRI and CT-imaging as intra-vitam diagnostic methods. Data from eight ball pythons including five spider ball pythons and three wild type ball pythons was assessed and evaluated comparatively. We were able to identify distinctive structural differences in inner ear morphology in spider ball pythons highly probable to relate to the wobble syndrome. To our knowledge, these anomalies are described for the first time and represent a basis for further anatomical and genetic studies and discussions regarding animal welfare in reptile breeding.

## Introduction

Reptile husbandry and breeding are established hobbies with a growing number of followers, playing an increasing role in the global pet industry. The US reptile market and industry - worth $1.4 billion - includes 4.7 million households keeping reptiles, 13.6 million reptiles kept as pets and 11.3 million reptiles bred / caught and exported each year (Collins and Fenili 2011). Within the European Union, exclusive the Baltic States, 7.9 million reptiles were kept as pets in 2019 (FEDIAF 2020). In the context of reptile breeding, morphs become increasingly popular among private owners and breeders due to their extraordinary looks and the profit margin achievable with rare individuals or new morphs. In this context, artificially selected morphs must be distinguished from morphs, which have occurred as consequence of natural or sexual selection (Andrèn and Nilson 1981; Calsbeek et al. 2010; Bastiaans et al. 2014; Fernández et al. 2017). Several thousand morphs of different species are available at present. Breeding efforts focus on new or brighter colours, new colour patterns and new variants of orientation, size and presence of scales.

Artificial selection and management in isolated populations is always associated with low individual numbers and many selection lines have experienced considerable inbreeding and loss of genetic variance. Laboratory animals, pets, cattle and horses have been studied extensively after occurrence of inbreeding depression and dissemination of many inherited defects, attributed to the extensive use of few preferred males (Leroy 2011). Defects can be phenotypically evident as, e.g., skeletal deformities or affect vigour, performance or fertility. To this point scientific examination of the effects of inbreeding and artificial selection in reptiles is rare but a plethora of studies was performed regarding other vertebrates. It was shown that the development of the skeletal system, especially the shape of the skull, of *Rattus villosissimus* was closely related to inbreeding (Lacy and Horner 1996). Similar effects on the skeletal system were noticeable in rabbits of different inbreeding levels Tanchev et al. 2011). Furthermore, a negative effect on flying performance of racing pigeons represented in the reduction of the median of the pigeon’s homing ability in races by 21% was shown and related to inbreeding (Meleg et al. 2005). Moreover, inbreeding was a good predictor for scute anomalies of European pond turtles (*Emys orbicularis*) observed in a study by Velo-Antón et al. (2011).

In addition, different associations between albinism or pigment alterations and anomalies of the inner ear or ability to hear have been shown in several species. Mair (1973) and Mair (1976) showed a correlation between coat colour, hair length and iris colour and hereditary deafness in white cats, and described cochlea-saccular degeneration in the dalmatian dog with hereditary deafness. Furthermore, auditory brainstem anomalies were described in albino cats (Creel et al. 1983), reduced neuronal size and dendritic length in the medial superior nucleus of albino rabbits (Conlee et al. 1986) together with differences in the endocochlear potential between albino and pigmented guinea pigs (Conlee and Bennett 1993). Moreover Ni et al. (2013) showed profound hearing deficit, characterized by elevated ABR thresholds, reduced distortion product otoacoustic emissions, absent endocochlear potential, loss of outer hair cells, and stria vascularis abnormalities correlating with embryonically fewer melanoblasts in mice with MITF mutations

With regards to morph breeding in reptiles, ball pythons (*Python regius*) are the most popular morphed reptiles. Currently 321 basic morphs (mostly assumed monogenetic) and more than 7300 designer morphs (combinations of different basic morphs) are confirmed and available (http://www.worldofballpythons.com/morphs/? 23.02.2022 3:30). As with other reptiles, ball python breeding is aimed at the production of different colours, patterns and scale anomalies. Several morphs, however, are reported to be associated with minor or major health issues, for example exophthalmos, reduced fertility, wobble syndrome or lethality (for reported morphs with their corresponding health issues see Table 1). Scientific validation for these disorders is scarce, but amongst breeders these anomalies are known, as they occur with distinct probabilities.

**Table 1:**
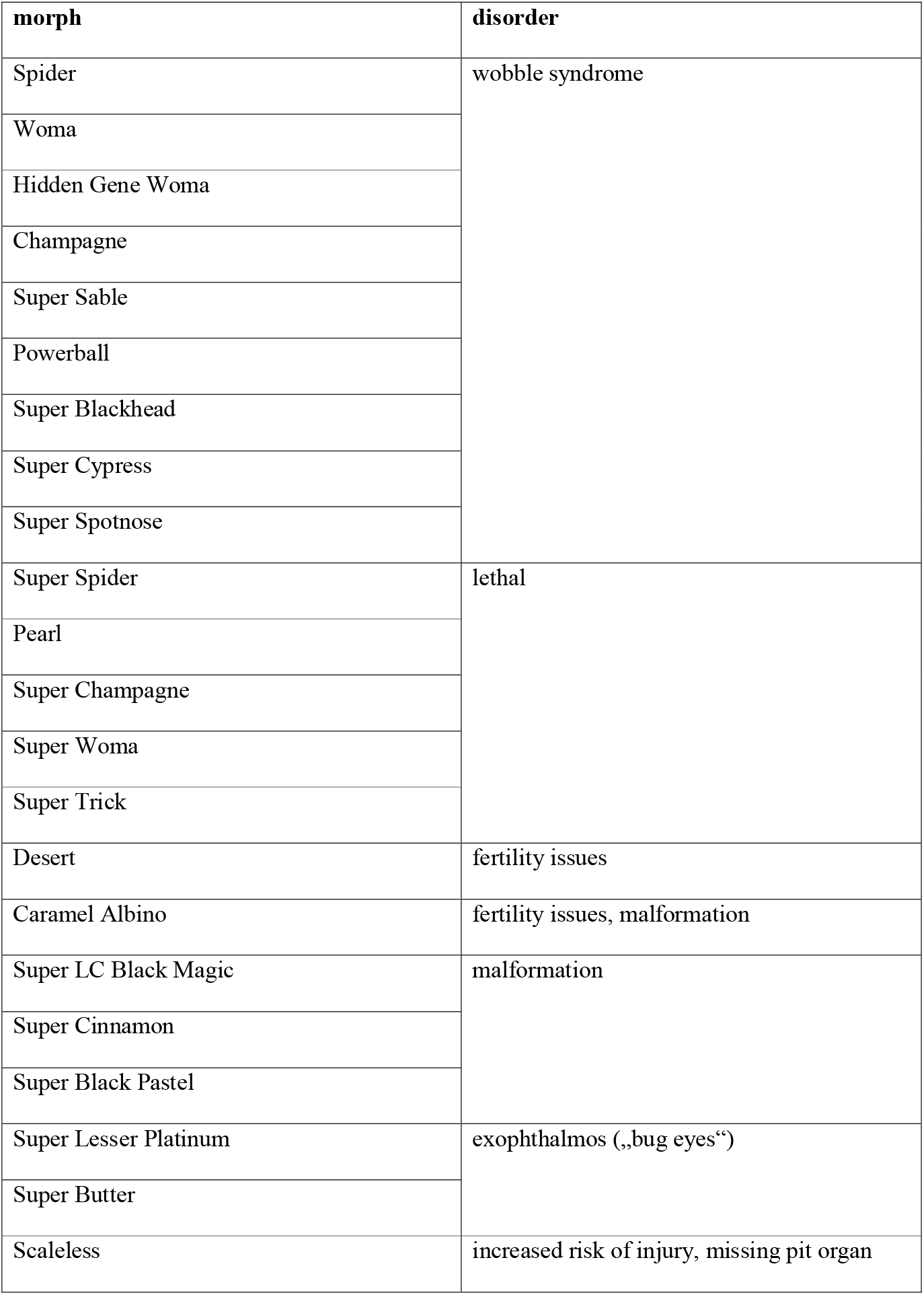
Known disorders and health issues of ball python morphs as reported on relevant websites and breeding literature. References: (Broghammer 2018, McCurley 2007, Öfner et al. 2018)

The wobbler syndrome is a collective term for clinical symptoms caused by a variety of neurological disorders in various species. It is a well-known disease in dogs (Mason 1979; da Costa 2010) and horses (Montali et al. 1974; Spaul et al. 1980). In dogs, the wobbler syndrome (caudal cervical spondylomyelopathy), is caused by different pathological changes resulting in compression of the caudal cervical spinal cord and subsequent neuronal damage. In contrast, in horses the wobbler syndrome is caused by cranial cervical vertebral myelopathy, a congenital condition that causes inflammation and arthritis in the cervical joints (Rooney 1972; Montali et al. 1974; Spaul et al. 1980). In both species, common symptoms include generalized weakness and ataxia and/or front limb lameness. In an analogous manner, symptoms reported in ball pythons and attributed to a wobbler-like syndrome involve head tremor, head tilting and twisting, reduced righting reflex and a lower accuracy at feeding (Rose and Williams 2014; Broghammer 2018).

Prior to this study Rose and Williams (2014) established a description of the wobble syndrome in spider ball pythons. The prevalence and the proportion of manifestation of the wobble syndrome were assessed through a survey of breeders and hobbyists. Furthermore, an animal welfare assessment was given by an expert group consisting of veterinarians and animal welfare scientists, who promoted a moderate to high impact on welfare in spider ball pythons.

Scientific information on the specific cause of the wobble syndrome is scarce and the association with morph breeding in ball pythons is correlational. In general, CNS symptoms can be caused by a variety of diseases in ball pythons. Viral infections such as Arenavirus (Inclusion Body Disease) (Chang and Jacobson 2010), Ferlavirus, Reovirus, Sunshine-Virus and Adenovirus (Marschang 2011) can cause symptoms like those of the wobble syndrome. Furthermore, bacterial and metabolic disorders can cause CNS symptoms if neural tissue is affected (Mader 2006). Anatomical anomalies caused by breeding deficiencies (e.g., high / low temperature; high / low moisture) or on a genetic basis seem to be another possible explanation. Pees (2015) suspected a congenital stenosis of the cervical spinal canal as the cause of the wobble syndrome in ball pythons but did not provide evidence to support this idea.

Medical advanced imaging techniques such as MRI and CT proved to be appropriate methods for intra vitam assessment of a variety of anatomical structures and anomalies in reptiles (Starck et al. 2015; Glodek et al. 2016) though the assessment of small structures may be subject to restrictions caused by imaging resolution. Considering the suggestion of a congenital anatomical anomaly (Pees 2015), the aim of this study was to determine, whether a morphological anomaly can be found and brought in association with the clinical occurrence of the wobble syndrome in spider ball pythons, using MRI and CT imaging as intra vitam diagnostic approaches.

## Materials and Methods

### Study Animals

Eight adult ball pythons were included into this study. Five “Spider” morphs were presented to the clinic by private owners to determine the cause of apparent neurological symptoms. Three ball pythons with natural phenotype (wild type) were used as comparison animals and underwent similar examination requested by a private owner precautionary to breeding. Animals were housed individually or in groups in terrariums (sizes at least 120 × 60 × 60 cm) with water ad libitum and shelter. Husbandry conditions were according to recommended standards (i.e., temperature gradient between 25°C and 30°C, 70 - 80% air humidity and an approximately 12 h light to 12 h dark regime) in all animals. Small deviations regarding temperature and humidity within the housing period were not found but cannot be excluded. Table 2 gives further information on age, sex and body mass of the animals. The study was conducted in accordance with the faculties ethical committee approval (VMF EK 4/2021).

**Table 2:**
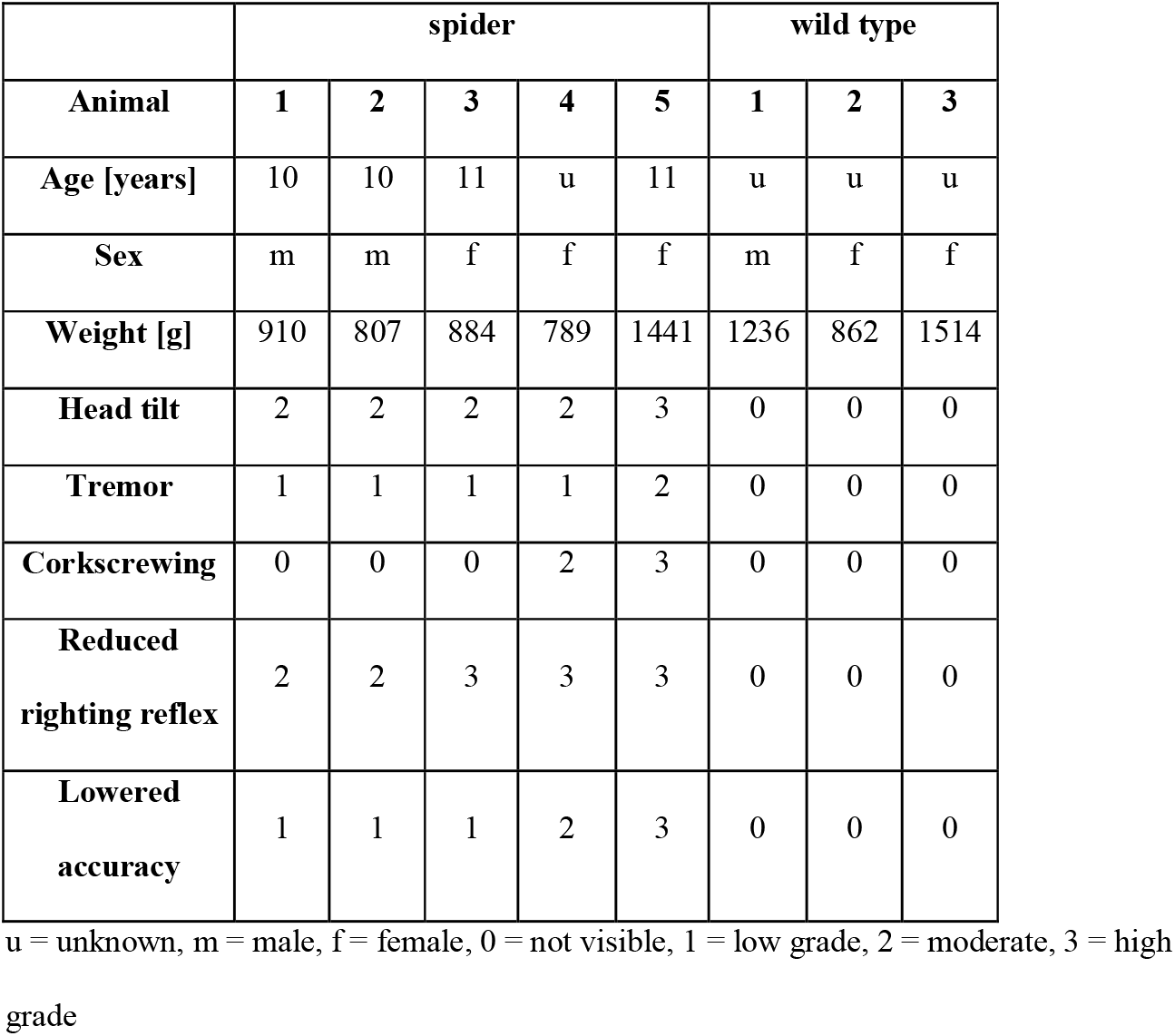
Clinical data on individuals used in the study.

### Clinical Examination and Microbiological Analysis

All snakes underwent a clinical examination including a neurological examination. The neurological examination was performed by two experienced reptile veterinarians and a specialist for small animal neurology (TF, DipECVN) and included observation regarding the mental status, behaviour and presence of any abnormal postures, as well as regularity of motion. Furthermore, the animals were tested for righting reflex and accuracy in striking prey (neurologic examination according to (Mader 2006) and deviations from normal state were scored according to the following system: 0 = not visible, 1 = low grade, 2 = moderate, 3 = high grade (see Table3 for further information). In one case of slight stomatitis choanal swabs, tracheal wash samples and cloacal swabs were examined for Adeno-, Arena-, Ferla-, Sunshine- and Reovirus, as well as bacterial and mycotic infection (aerobic bacterial culture on columbia sheep blood agar, mac conkey no 3 and brilliant green bile agar and mycological culture on sabouraud medium; incubation over 48 hours for bacteria and up to 7 days for fungi at 30°C).

**Table3:** Scoring of symptoms in the clinical and neurological examination of the ball pythons.

| Grade | Visible at rest | Visible after<br><1min handling | Visible after<br>>1min handling | Visible while<br>feeding |
| --- | --- | --- | --- | --- |
| 1 = Not visible | no | no | no | no |
| 2 = Low | no | no | yes | yes |
| 3 = Moderate | no | yes | yes | yes |
| 4 = High | yes | yes | yes | yes |

### Computed Tomography and Magnetic Resonance Imaging

The snakes were anaesthetized with ketamine (20 mg/kg i.m.) and medetomidine (0.2 mg/kg i.m.) (Pees 2015) and examined in a normal stretched ventral position using a six-slice spiral CT scanner (Brilliance CT 6; Philips, Hamburg, Germany) and an MRI scanner (Ingenia 3T, Philips, Hamburg, Germany). Transversal contiguous CT-slices, as well as sagittal, transversal and dorsal MRI-scans (T1- and T2-weighted) were obtained by scanning an area from the nostril to about 15 % of body length. The CT settings were: collimation 0.75 mm, slice thickness 1.0 mm, pitch 0.6, with adaptation of the helix for an increment of 0.5 mm, start at tube voltage 120 kV and 120 mA with adaptation of current intensity.

Objective of the CT scans was to assess the bone structure of the skull and the vertebral canal regarding uniformity, bilateral symmetry and width of the cranial and tympanic cavities (Boistel et al. 2011; Krautwald-Junghans et al. 2011; Palci et al. 2017).

Objective of the MRI scans was to assess neuronal tissue of the brain and the spinal cord, and evaluation of peri- and endolymphatic spaces. T2 images in the transversal and dorsal plane were selected for further assessment. Evaluation and measurements were performed using the software IntelliSpace^®^ PACS Enterprise 4.4.543.0 (Phillips, Hamburg) and measuring points and areas were chosen and marked manually. In dorsal images, the vertebral canal was examined regarding consistent width and presence of osseous stenosis. Furthermore, bilateral symmetry and homogeneity of brain tissue (Northcutt 2013; Naumann et al. 2015; Allemand et al. 2017), spinal cord, peripheral neurons, if evaluable, and the inner ear were determined, as well as the dimension of the bullae and the semi-circular canals was evaluated on both planes.

### Measurements

Based on the MRI scans, measurements were taken on both sides of each individual snake to determine the size of the inner structures in relation to each other as well as the body size of the snake. The bony labyrinth i.e., the peri- and endolymphatic space inside the prootic bone is one possible sources of the symptoms. Therefore, we measured the area within the inner diameter of the semicircular canals at its greatest extents in transversal and dorsal planes. As reference points, we measured the vitreous body of the eye at its greatest extent, the diameter of the telencephalon and the medulla caudal to the peri- and endolymphatic space of the inner ear at their greatest widths in transversal and dorsal planes. We then calculated ratios of the bony labyrinth area to the diameter of the telencephalon, the medulla and the vitreous body. These were computed to compensate for size differences between study animals and compared between wild type and spider ball pythons. The images of the three wild type ball pythons, as well as publications (Boistel et al. 2011; Christensen et al. 2012; Palci et al. 2017) were consulted as anatomical reference to the five spider ball pythons.

### Statistical Analysis

To account for size differences between individuals, proportions of the inner ear to the telencephalon and the vitreous body were calculated and used for further statistical analysis. Proportions were compared between left and right side within the groups of spider and wild type ball pythons as well as between spider and wild type ball pythons.

Statistical analysis was performed using the program SPSS 25.0 (IBM, Armonk, USA). Based on the low number of samples and confirmed by the Kolmogorov-Smirnov-Test, no standard distribution was assumed for the data obtained. Therefore, the Mann-Whitney-U-test was used to determine significance of differences between both groups. Significance was assumed with p ≤ 0.05.

## Results

### Clinical Examination and Microbiological Analysis

The wild type ball pythons were healthy on clinical and neurological examination. The spider ball pythons, except for one individual showing minor signs of stomatitis, displayed no signs of any disease apart from the neurological deficits described below. Choanal swabs, tracheal wash samples and cloacal swabs from the snake with stomatitis proved to be negative for Adeno-, Arena-, Ferla-, Sunshine- and Reovirus (PCR detection), as well as unremarkable in the bacteriological (mixed culture including Enterobacteriaceae, Pseudomonadaceae) and mycological examination (assessment of cultured bacteria according to (Résière et al. 2018; Hill et all. 2018; Tang et al. 2018; Zhang et al. 2019), no fungi detected).

### Neurological examination

The neurological examination in the wild type group was completely unremarkable. In the Spider morph group, neurological symptoms ranged from head tilt (moderate in four animals, high grade in one animal), tremor (low grade to moderate in all five animals), corkscrewing (moderate in one animal and high grade in one animal), reduced striking accuracy (moderate and high grade in one animal each, low grade in the remaining three) to reduced righting reflex (moderate in two animals, high grade in three animals) (detailed results see Table 2).

### Computed Tomography

In the wild type ball pythons, the vertebral canal appeared consistent in width and shape with no evidence of osseous protrusions or stenosis. Furthermore, no evidence of vertebral or vertebral joint malformation was observed (see Figure 1). The atlanto-occipital joint was symmetric, the articular surfaces were even, and no anomalies were observed. The wedge-shaped skull expanded laterally to reach a maximum width at the level of the otic capsules. The neurocranium formed a bilateral symmetric and regular shell with steady bone thickness congruent to prior studies (Boistel et al. 2011; Christensen et al. 2012; Palci et al. 2017). The posterior margin of the orbits was located halfway between rostral tip of the skull and the atlanto-occipital joint. The orbit formed consistent round structures parallel to the ocular bulb rostro-lateral to the cranial cave. Circumscribed osseous formations within the prootic bone showed central oval osseous structures surrounded by semi-radiolucent areas confined by the outer skull and the osseous boundary of the cranial cavity (see Figure 2).

**Figure 1:**
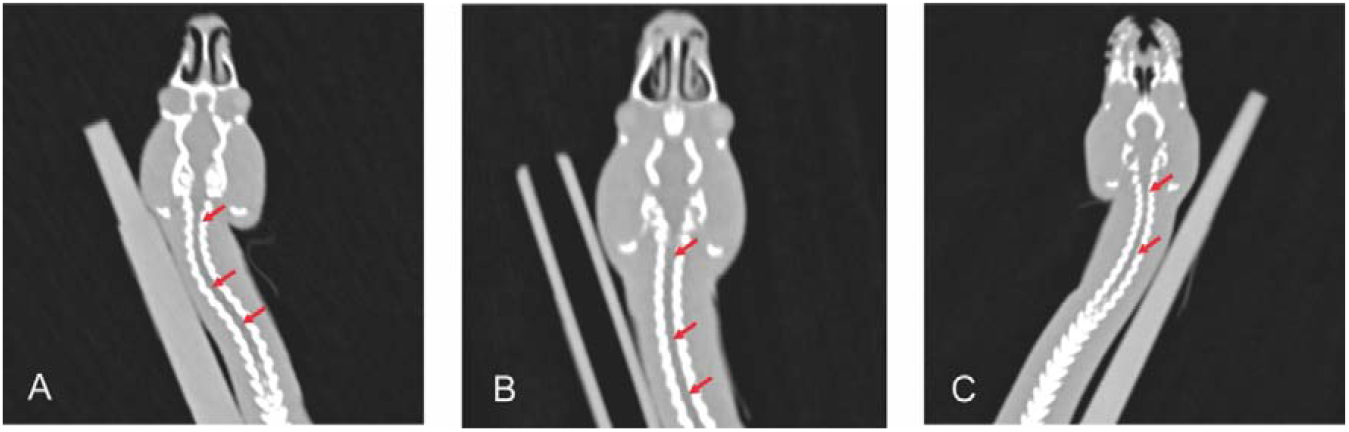
CT image, dorsal plane, cervical vertebral canal of three spider ball pythons. Arrows, Cervical vertebral canal.

**Figure 2:**
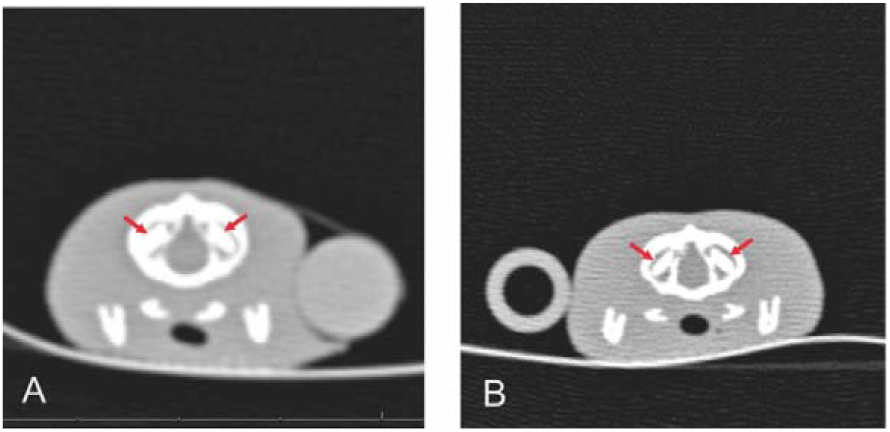
CT image, transversal plane. The inner ear of two wild type ball pythons. Arrows, central osseus structure in the area of the inner ear.

In all spider ball pythons, the anatomy of the vertebral canal and the skull were consistent with the findings in wild type animals. The presentation of the inner ear and the bony labyrinth structure, however, showed differences to the wild type ball pythons in shape and manifestation of the central oval osseous structure. In two animals (Spider 4 and Spider 5; see Table 2 for further information on animals), the area within the osseous boundaries was filled with semi-radiotranslucent material and the radiodense central structure was non-existent (see Figure 3).

**Figure 3:**
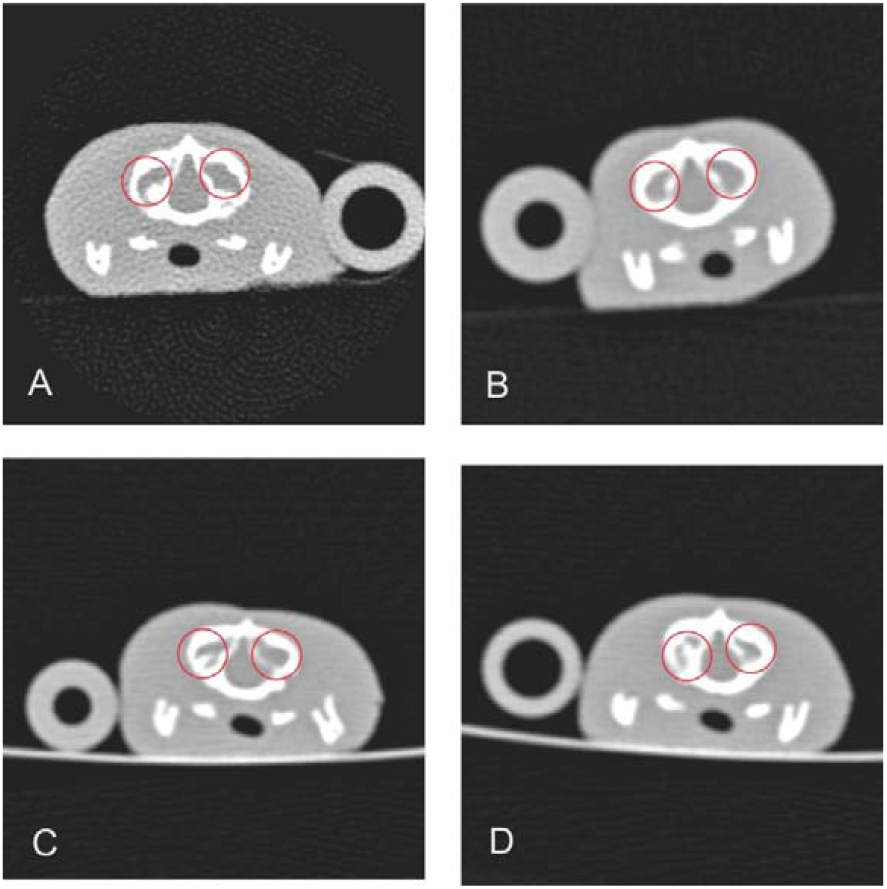
CT image of the inner ear of four spider ball pythons (transversal plane). Circles, area of the inner ear.

### Magnetic Resonance Imaging

In wild type ball pythons, the neuronal tissue of the brain and the spinal cord as well as the peripheral nervous system was shown as medium intense signal in T2 images with distinct more and less intense bilateral symmetric structural features. No spinal disk protrusions or other structural abnormalities were identified (see Figure 4). Wild type ball pythons exhibited bilateral symmetric intense signal semicircular areas with a median diameter of 0.7 mm (range 0.7 to 0.8 mm) surrounding a central very low intensity signal structure rostrolateral of the medulla congruent to the central oval osseous structure as seen on the CT-images (see Figure 5) (Krautwald-Junghanns et all.2011). In spider ball pythons, the semicircular areas were non-symmetric and widened. The central low intensity signal structures were smaller and amorphous in one spider ball python (Spider 1; see Table 2 for further information on animals), non-existing on one side and reduced in size on the other in two cases (Spider 2 and Spider 3; see Table 2 for further information on animals) and non-existent on both sides in two spider ball pythons (Spider 4 and Spider 5; see Table 2 for further information on animals). The low intensity signal structures were replaced by high intensity signal filled areas in all cases (see Figure 6). Congruent to the CT imaging findings, the vertebral canal, as well as the cranial cavity did not show any malformations and appeared equivalent to the comparison animals.

**Figure 4:**
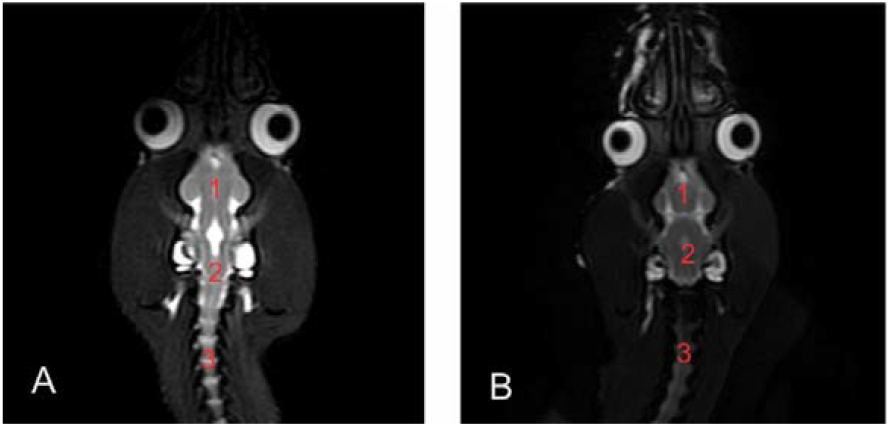
T1 MRI images of the brain and cervical spinal cord of two spider ball pythons. 1, telencephalon; 2, medulla; 3 spinal cord

**Figure 5:**
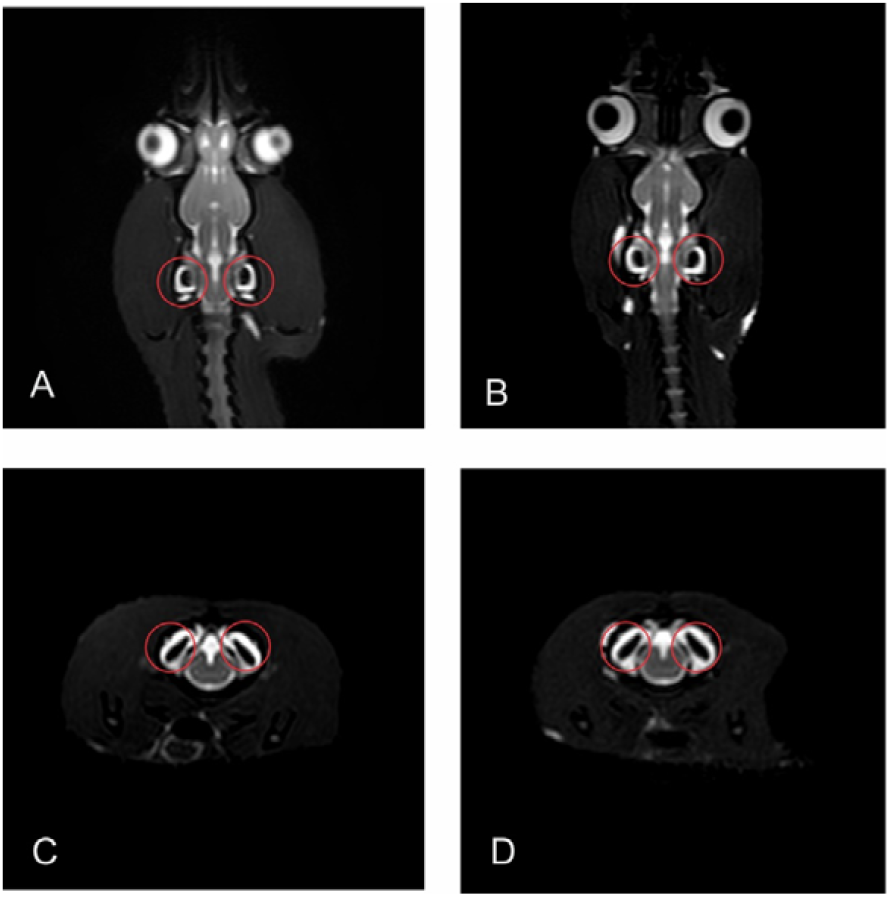
T1 MR images of the inner ear of two wild type ball pythons. A and B transversal plane. C and D dorsal plane. Circles, area of the inner ear.

**Figure 6:**
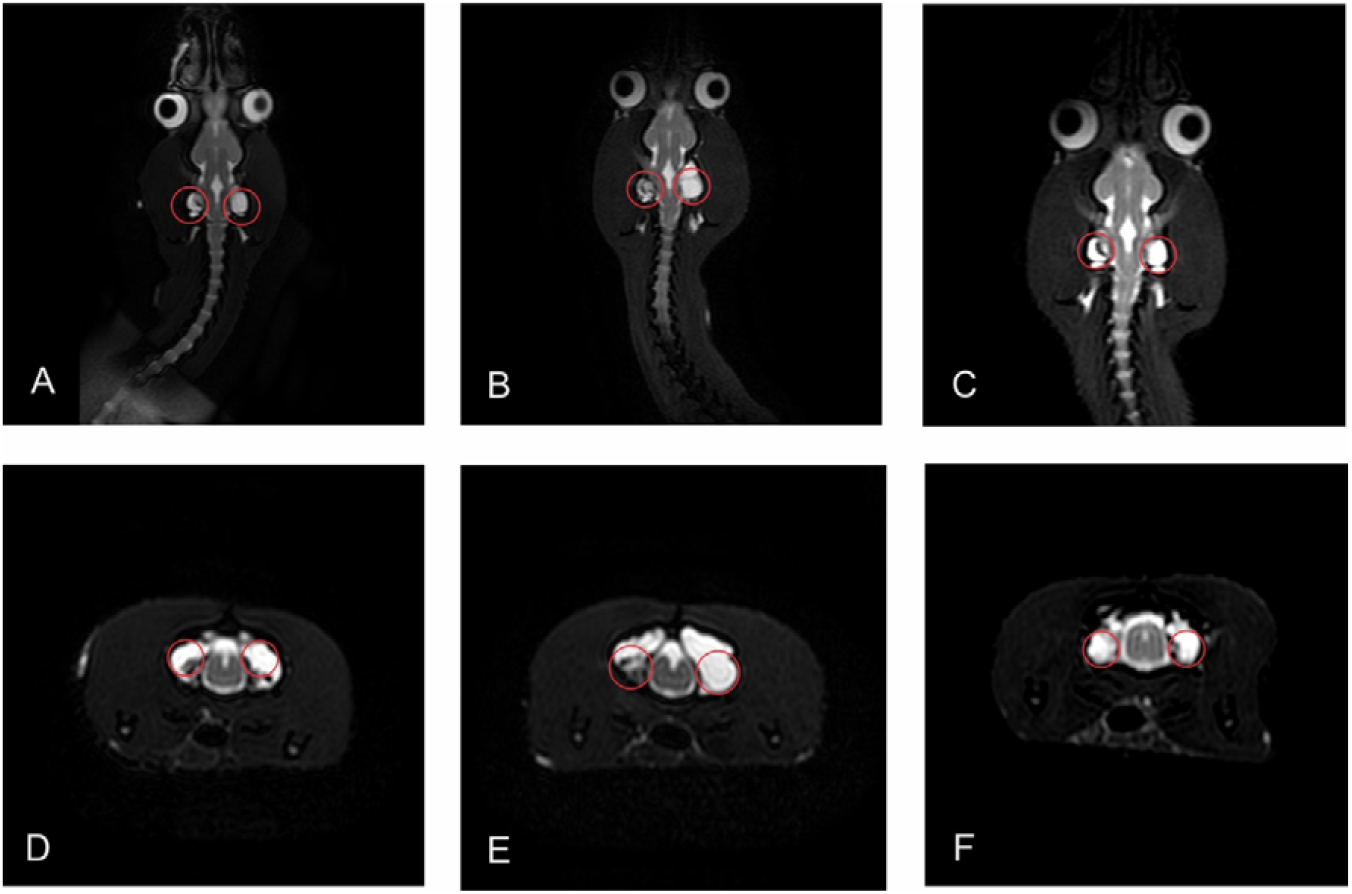
T1 MR images of the inner ear of three spider ball pythons. A, B and C dorsal plane. D, E and F transversal plane. Circles, area of the inner ear.

### Measurements

In wild type ball pythons, the median width of the telencephalon was 8.95 mm (range from 8.70 to 9.10 mm), the median medulla width was 4.55 mm (range from 3.90 to 4.90 mm) and the median vitreous body dimension was 8.75 mm^2^ (range from 7.38 to 11.13 mm^2^). In spider ball pythons, median width of the telencephalon was 8.70 mm (range from 8.20 to 9.10 mm), the median medulla width was 4.50 mm (range from 3.90 to 5.10 mm) and the median vitreous body dimension was 8.34 mm2 (range from 6.24 to 10.77 mm2). The two groups did not differ significantly regarding these data. The median dimension of the inner ear, however, differed significantly (p < 0.05) between spider ball pythons with 0 mm2 (range from 0 to 1.21 mm2) and wild type ball pythons with 2.77 mm2 (range from 2.24 to 3.12 mm2). For detailed information regarding measurements see Table 4.

**Table 4:**
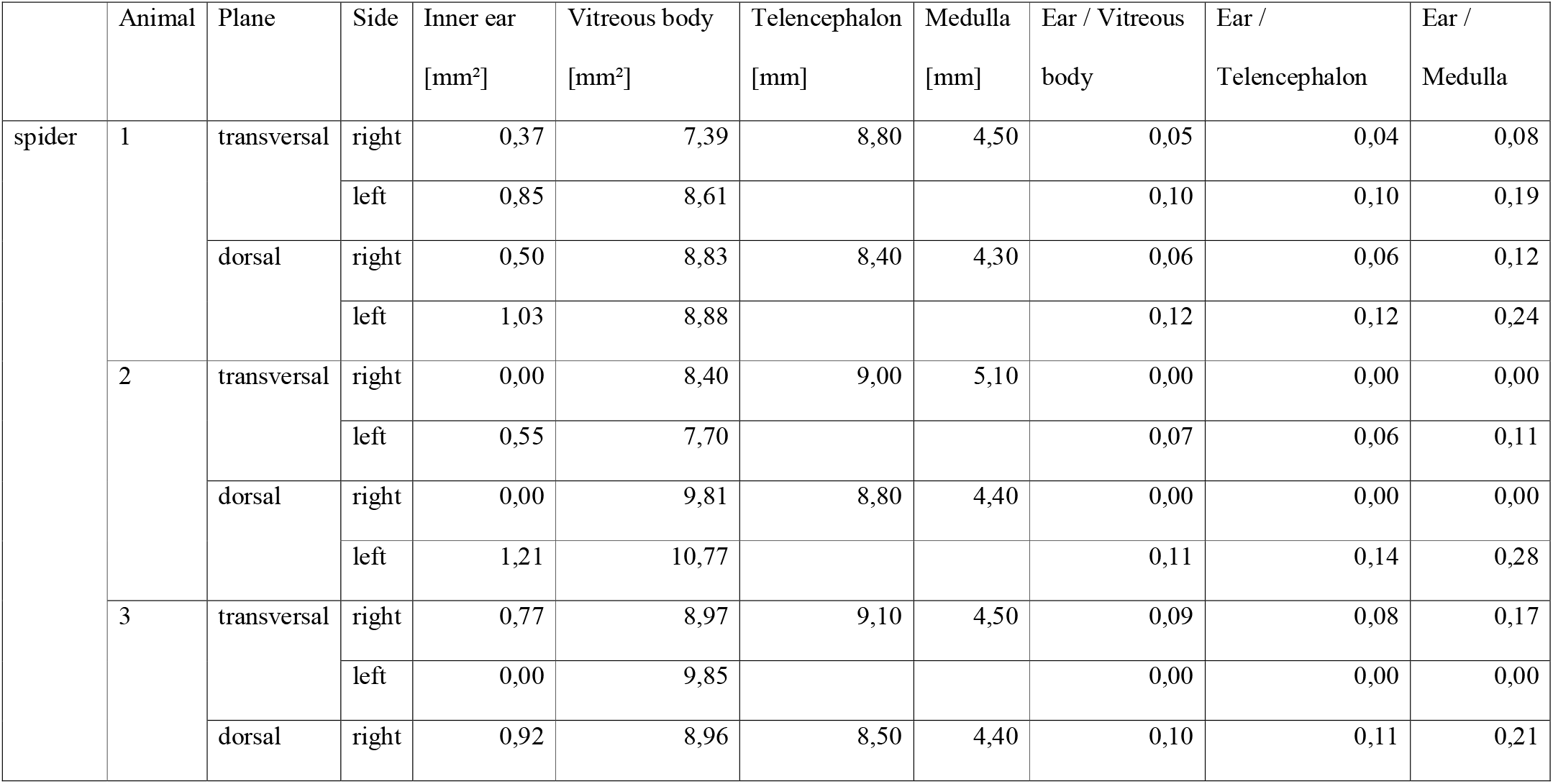
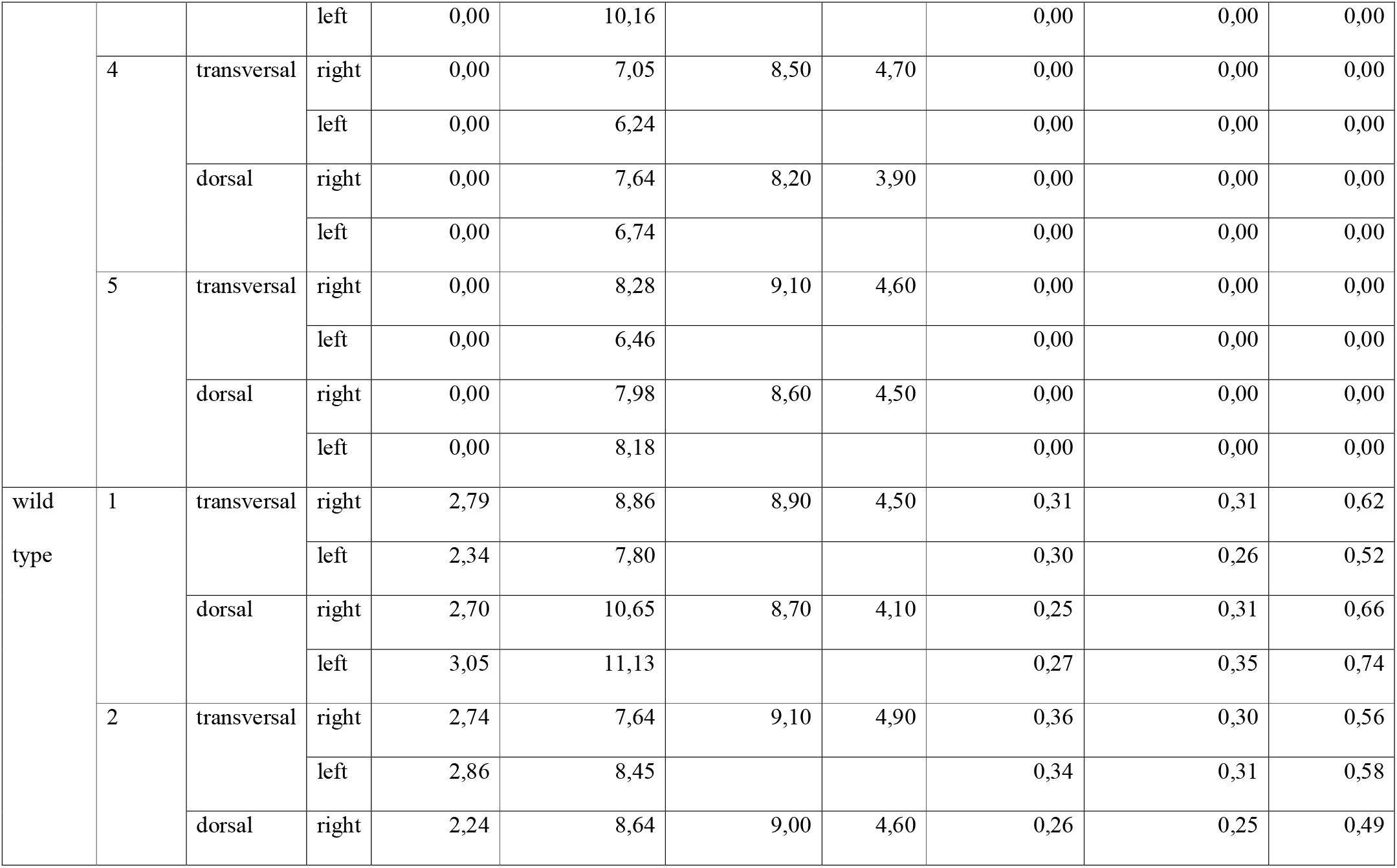
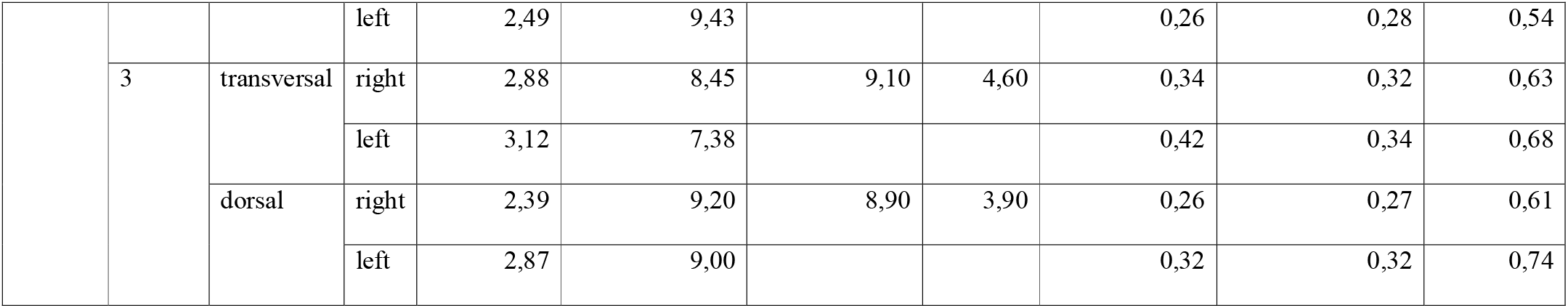
Measurements of the dimension of the inner ear, the dimension of the vitreous body, the width of the telencephalon and the width of the medulla of the ball pythons within MR images. Proportions of the inner ear to the vitreous body, the telencephalon and the medulla.

In wild type ball pythons, proportions of inner ear dimension to width of the telencephalon ranged between 0.25 and 0.35 (median 0.31), inner ear dimension to medulla width ranged between 0.49 and 0.74 (median 0.62) and inner ear dimension to vitreous body dimension ranged between 0.25 and 0.42 (median 0.31). In contrast, the proportion of the inner ear to the width of the telencephalon and medulla and the dimension of the vitreous body in spider ball pythons ranged between 0.00 and 0.14 (median 0.00), 0.00 and 0.28 (median 0.00) and 0.00 and 0.12 (median 0.00) respectively. The Mann-Whitney-U-test showed a significant difference between the wild type and spider ball pythons regarding proportion of the inner ear compared to width of the telencephalon and medulla and the dimension of the vitreous bodies (p < 0.05) as shown in Figures 7, 8 and 9.

**Figure 7:**
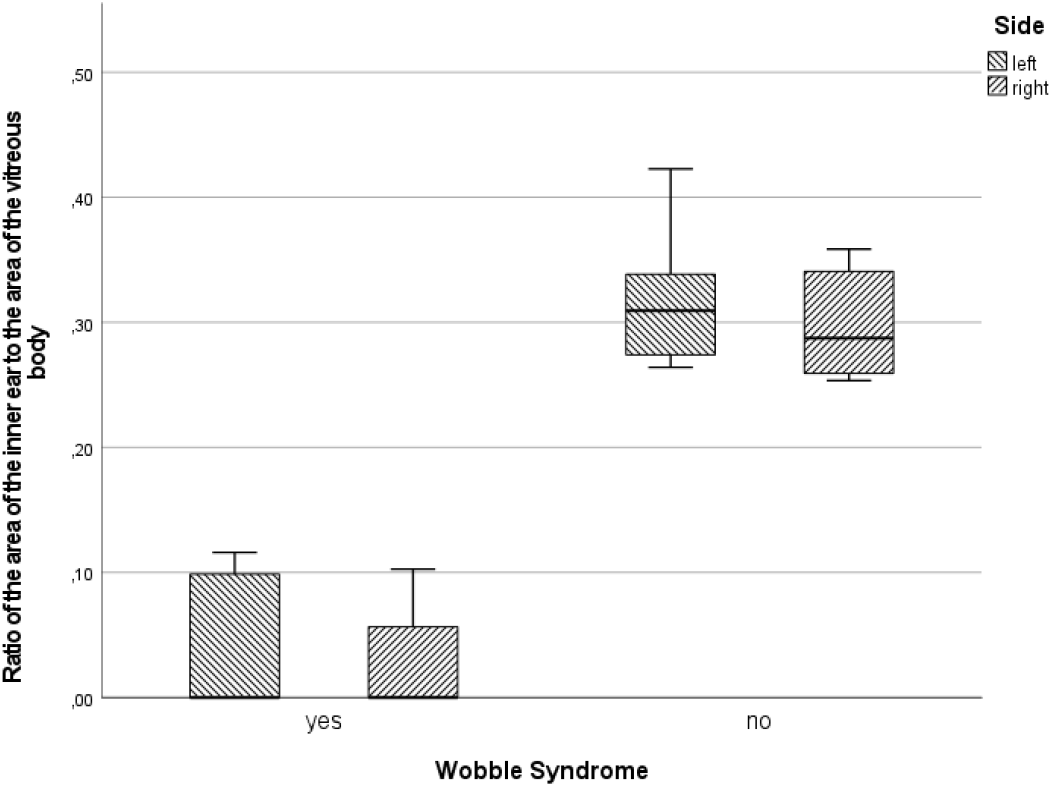
Boxplot showing the proportion of the area of the inner ear to the vitreous body comparing spider (with wobble syndrome) and wild type ball pythons.

**Figure 8:**
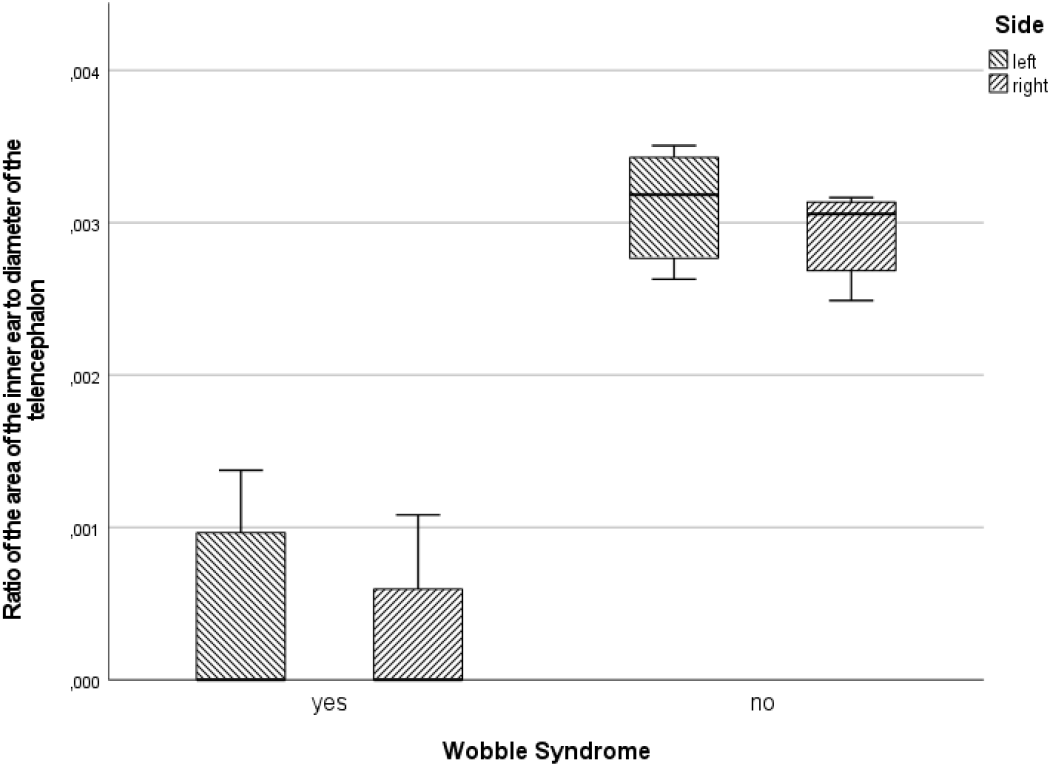
Boxplot showing the proportion of the area of the inner ear to the width of the telencephalon comparing spider (with wobble syndrome) and wild type ball pythons.

**Figure 9:**
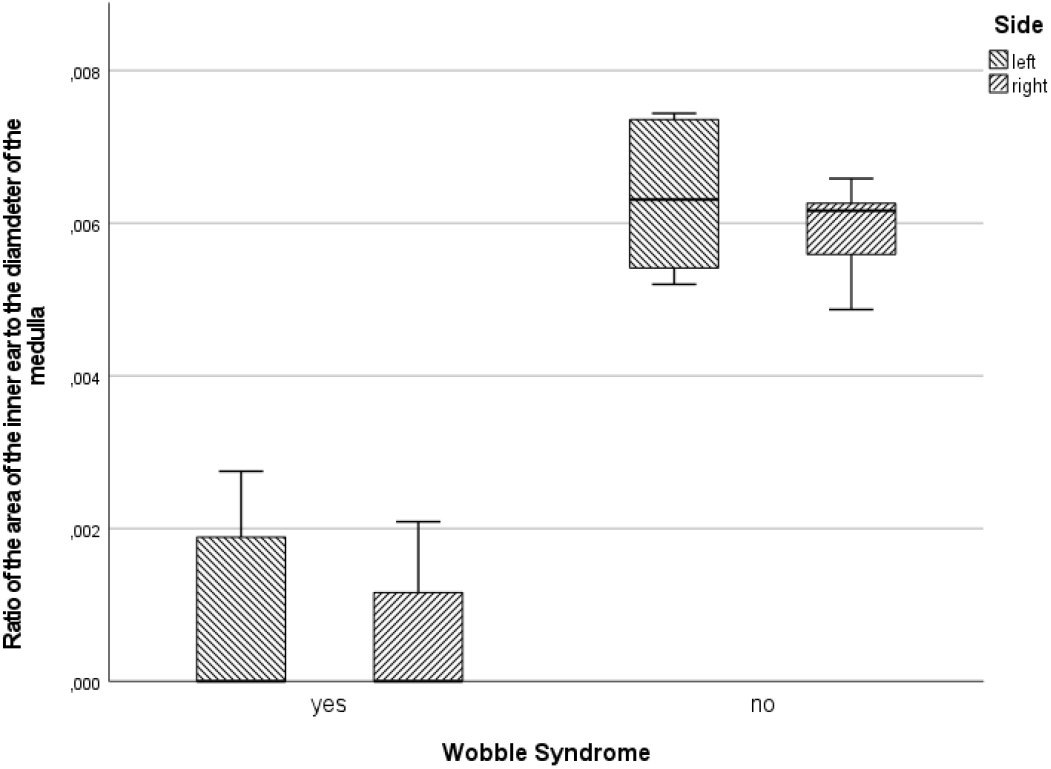
Boxplot showing the proportion of the area of the inner ear to the width of the medulla comparing spider (with wobble syndrome) and wild type ball pythons.

## Discussion

The present study was designed to find out if specific morphological variations are associated with the wobble syndrome in the spider morph of the ball python. To the best of our knowledge, there has been no similar study investigating morph associated diseases in spider ball pythons except for Rose and Williams (2014).

The neurological symptoms described by Rose and Williams (2014), namely tremor, opisthotonos, “corkscrewing”, reduced accuracy and delayed or missing righting reflex, were confirmed during clinical examination in all five spider ball pythons to varying degrees (see Table 2), but were absent in all wild type animals examined. In all wild type pythons osseous and neural structures were considered unremarkable in accordance with current literature, while all spider morph ball pythons showed abnormal structures of the osseous and neural parts of the inner ear; despite the relatively low spatial resolution we achieved in this study.

Our clinical examination scientifically confirmed what has been common knowledge among breeders and owners for a considerable period and added various information and details concerning the morphological basis of the wobble syndrome. The combined use of computed tomography and magnetic resonance imaging proved to be an appropriate method to examine the osseous and non-osseous structures of the head and cranial vertebral region of ball pythons and showed congruency with anatomical descriptions (Maisano and Rieppel 2007; Rieppel et al. 2009; Olorie 2010). Furthermore, we were able to relate to the results of Palci et al. (2017) and Christensen et al. (2012) regarding the morphology of the inner ear in healthy ball pythons.

We had expected to find cervical spinal lesions affecting the spinal cord resulting in similar clinical symptoms as described for the wobbler syndrome in dogs and horses, but we did not identify any spinal pathology. In contrast, we found obvious morphological deviations in the inner ear of the spider ball pythons. Those changes are most likely interfering with the function of the receptor organ of the vestibular system located in the inner ear. The abnormalities seen on CT and MR imaging are probably causing typical symptoms of asymmetric vestibular input to the vestibular nuclei of the brainstem such as head tilt and corkscrewing. Even the reduced accuracy of catching prey in the spider python may be a manifestation of a loss of spatial orientation caused by vestibular system malfunction.

However, the exact mechanism of a potential causative relationship between the phenotypical appearance of the spider ball python and the pathological changes of the inner ear seen on CT and MR images has still to be determined.

Prior to this study, several cases of hereditary deafness or inner ear malformation additionally to other pleiotropic effects have been investigated and linkage between these and pigmentation genes has been established. Bellone (2010) summarized six such pleiotropic effects in horses, which include neurological defects (lethal white foal syndrome and lavender foal syndrome), hearing defects, eye disorders (congenital stationary night blindness and multiple congenital ocular anomalies), as well as horse-specific melanoma. Furthermore, it was shown that hereditary hearing defects in deaf white cats affect both sensory epithelium and neural structures (Pujol et al. 1977). In humans, Waardenburg (1951) described a syndrome combining developmental anomalies of the eyelids, eyebrows and nose root with pigmentary defects of the iris and head hair and with congenital deafness. This study was further affirmed by Fisch (1959) who showed additional symptoms and a distinct characteristic of partial hearing loss in patients (better hearing for higher frequencies). The Tietz syndrome, which includes congenital profound deafness and generalised hypopigmentation, was shown to be caused by mutation of the MITF gene. Mutations in other regions of this gene produce Waardenburg syndrome type 2 (Smith et al. 2000). Additionally, Dawoud et al. (2017) showed that 50% of vitiligo patients suffered from peripheral vestibular disorders in addition to auditory affection. In this study, hearing defects were not assessed due to the nature of sound perception in snakes through mainly bone and body conducted vibration and difficult assessment of reaction to sound but should be considered as an examination tool in addition to medical imaging in follow up studies.

Breeders attempting to outbreed the wobble syndrome while maintaining the specific pattern failed so far, which implies a strong link between the wobble syndrome and the spider pattern.

Breeders mostly believe the spider mutation to be autosomal dominant. The inheritance and the exact genetic basis of the spider morph and the wobble syndrome should be evaluated to further deepen the understanding of morph associated diseases. The occurrence of the wobble syndrome in several different morphs with reduced dark pattern leads to the assumption that an alteration of melanocyte and pigmentation development may cause the morphological anomalies, which manifest as malformation of the inner ear in spider ball pythons. The morphological and genetic basis for the syndrome in other morphs is yet unknown and needs to be investigated. Embryologically, the ectoderm and the neural crest figure prominently in the development of several tissues ranging from neurons and glial cells of the peripheral nervous system to pigment cells, fibroblasts to smooth muscle cells, and odontoblasts to adipocytes (Mayor and Theveneau 2013). The inner ear derives almost completely from cells of the otic placode, which is part of the ectoderm (Whitfield 2015). Mutations affecting pigmentation are therefore likely to have pleiotropic effects throughout the organism in humans, mammals and vertebrates in general and especially regarding the inner ear.

Since all five spider ball pythons included in this study exhibited the wobble syndrome in differing degree and specific symptoms and showed similar morphological anomalies, namely malformation of the saccule (Starck et al. submitted) and the inner ear, it is highly probable that these alterations are the cause of the wobble syndrome. These findings need to be further verified by larger sample size and of high-resolution imaging like μ-CT and / or histology. Furthermore, specific genetic studies are needed to confirm and describe the genetic defect that could possibly lead to the found anomalies. Moreover, only macroscopic morphological findings in this study were validated as structural differences in spider ball pythons. Nonetheless this study documents that inner ear malformations in spider ball pythons may be detected in live animals with comparatively crude but easily accessible clinical imaging. More detailed information may be obtained by means of μ-CT but are impractical for diagnostics in clinical routine due to its limitation to postmortem assessment.

In terms of animal welfare this study gives a foundation for further discussion and research but should be complemented with additional studies dealing with a wide variety of morphs of different species and their genetic basis.

Referring to the occurrence of neurological symptoms, we assumed a moderate to high impact on welfare in these animals, as promoted by Rose and Williams (Rose and Williams 2014). Essential for our assumption was the inability of precisely catching prey and the reduced ability of orientation and balance.

As individuals that lack the specific pattern mutation do not express the wobble syndrome (Rose and Williams 2014) this could, in our opinion, justify prohibition of further breeding. Resting on the estimation of a high impact on welfare and the associated suffering in cases of high-grade symptoms and partial inability of natural behaviour euthanasia of highly affected individuals may be appropriate.

## References

Allemand R, Boistel R, Daghfous G, Blanchet Z, Cornette R et al. (2017) Comparative morphology of snake (Squamata) endocasts: evidence of phylogenetic and ecological signals. Journal of Anatomy, 231 (6), 849–868. DOI: 10.1111/joa.12692.

Anderson Maisano J, Rieppel O (2007) The skull of the Round Island boa, Casarea dussumieri Schlegel, based on high-resolution X-ray computed tomography. Journal of Morphology, 268 (5), 371–384. DOI: 10.1002/jmor.10519.

Andrén C, Nilson G (1981) Reproductive success and risk of predation in normal and melanistic colour morphs of the adder, Vipera berus. Biological Journal of the Linnean Society, 15 (3), 235–246. DOI: 10.1111/j.1095-8312.1981.tb00761.x.

Bastiaans E, Bastiaans M J, Morinaga G, Castañeda Gaytán J G, Marshall J C et al. (2014) Female preference for sympatric vs. allopatric male throat color morphs in the mesquite lizard (Sceloporus grammicus) species complex. PloS One, 9 (4), e93197. DOI: 10.1371/journal.pone.0093197.

Bellone R R (2010) Pleiotropic effects of pigmentation genes in horses. Animal Genetics 41 Suppl 2, 100–110. DOI: 10.1111/j.1365-2052.2010.02116.x.

Boistel R, Herrel A, Lebrun R, Daghfous G, Tafforeau P et al. (2011) Shake rattle and roll. The bony labyrinth and aerial descent in squamates. Integrative and Comparative Biology, 51 (6), 957–968. DOI: 10.1093/icb/icr034.

Broghammer S (2018) Python Regius. Atlas der Farbmorphen Pflege und Zucht. 2nd Edition. Natur und Tier Verlag, Münster

Calsbeek B, Hasselquist D, Clobert J (2010) Multivariate phenotypes and the potential for alternative phenotypic optima in wall lizard (Podarcis muralis) ventral colour morphs. Journal of Evolutionary Biology, 23 (6), 1138–1147. DOI: 10.1111/j.1420-9101.2010.01978.x.

Chang L, Jacobson E R (2010) Inclusion Body Disease, A Worldwide Infectious Disease of Boid Snakes: A Review. Journal of Exotic Pet Medicine, 19 (3), 216–225. DOI: 10.1053/j.jepm.2010.07.014.

Christensen Bech C, Christensen-Dalsgaard J, Brandt C, Teglberg Madsen P (2012) Hearing with an atympanic ear. Good vibration and poor sound-pressure detection in the royal python, Python regius. The Journal of Experimental Biology, 215 (Pt 2), 331–342. DOI: 10.1242/jeb.062539.

Collis A H, Fenili R N (2011) The modern US reptile industry. Washington, DC: Georgetown Economic Services.

Conlee J W, Bennett M L (1993) Turn-specific differences in the endochochlear potential between albino and pigmented guinea pig. Hearing Research, (65), 141–150.

Conlee J W, Parks T N, Creel D J (1986) Reduced Neuronal Size and Dendritic Length in Medial Superior Olivary Nucleus of Albino Rabbits. Brain Research, (363), 28–37.

Creel D, Conlee J W, Parks T N (1983) Auditory Brainstem Anomalies in Albino Cats. I. Evoked Potential Studies. Brain Research, (260), 1–9.

da Costa R C (2010) Cervical spondylomyelopathy (wobbler syndrome) in dogs. The Veterinary clinics of North America. Small Animal practice, 40 (5), 881–913. DOI: 10.1016/j.cvsm.2010.06.003.

Dawoud E A E, Ismail E I, Eltoukhy S A, El-Sharabasy A E (2017) Assessment of auditory and vestibular functions in vitiligo patients. Journal of Otology, 12 (3), 143–149. DOI: 10.1016/j.joto.2017.07.001.

FEDIAF (2020) European Facts & Figures 2019. FEDIAF. https://fediaf.org/images/FEDIAF_facts_and_figs_2019_cor-35-48.pdf, lastly checked 12.12.2021.

Fernández J B, Bastiaans E, Medina M, La Méndez De Cruz F R, Sinervo B R et al. (2017) Behavioral and physiological polymorphism in males of the austral lizard Liolaemus sarmientoi. Journal of Comparative Physiology. A, Neuroethology, Sensory, Neural, and Behavioural Physiology, 204(2):219–230. DOI: 10.1007/s00359-017-1233-1.

Fisch L (1959) Deafness as part of a hereditary syndrome. In: The Journal of Laryngology and Otology, 73, S. 355–382. DOI: 10.1017/s0022215100055420.

Glodek J, Adamiak Z, Przeworski A (2016) Magnetic resonance imaging of reptiles, rodents and lagomorphs for clinical diagnosis and animal research. Comparative Medicine, 66 (3), 216–219.

Hill A J, Leys J E, Bryan D, Erdman F M, Malone K S et al. (2018) Common Cutaneous Bacteria Isolated from Snakes Inhibit Growth of Ophidiomyces ophiodiicola. EcoHealth, 15 (1), S. 109–120. DOI: 10.1007/s10393-017-1289-y.

Krautwald-Junghanns M, Pees M, Reese S, Tully T (2011): Diagnostic Imaging of Exotic Pets. Birds - Small Mammals - Reptiles. Schlütersche Verlagsgesellschaft mbH & Co. KG. Hannover.

Lacy R C, Horner B E (1996) Effects of inbreeding on skeletal development of Rattus villosissimus. Journal of Heredity, 87 (4), 277–287.

Lattanzio M S, Metro K J, Miles D B (2014) Preference for male traits differ in two female morphs of the tree lizard, Urosaurus ornatus. PloS One, 9 (7), e101515. DOI: 10.1371/journal.pone.0101515.

Leroy G (2011) Genetic diversity, inbreeding and breeding practices in dogs: results from pedigree analyses. Veterinary Journal (London, England: 1997), 189 (2), 177–182. DOI: 10.1016/j.tvjl.2011.06.016.

Mader D R (2006): Reptile Medicine and Surgery. 2nd Edition. Saunders Elsevier. St. Louis.

Mair I W S (1973) Hereditary Deafness in the White Cat. Acta oto-laryngologica, 76 (sup314), 5–48. DOI: 10.1080/16512251.1973.11675740.

Mair I W S (1976): Hereditary deafness in the Dalmatian dog. Archives of Oto-Rhino-Laryngology, (212), 1–14.

Marschang R E (2011) Viruses infecting reptiles. Viruses, 3 (11), 2087–2126. DOI: 10.3390/v3112087.

Mason T A (1979) Cervical vertebral instability (wobbler syndrome) in the dog. The Veterinary Record, 104 (7), 142–145.

Mayor R, Theveneau E (2013) The neural crest. Development (Cambridge, England), 140 (11), 2247–2251. DOI: 10.1242/dev.091751.

McCurley (2007) Python Regius: Das Kompendium. Edition Chimaira, Frankfurt am Main.

Meleg I, Pakuts G, Reiczigel J (2005) Inbreeding effect on flying performance of racing pidgeons. Archiv für Geflügelkrankheiten, 69 (1), 23–26.

Montali R J, Bush M, Sauer R M, Gray C W, Xanten W A (1974) Spinal ataxia in zebras. Comparison with the wobbler syndrome of horses. Veterinary Pathology, 11 (1), 68–78. DOI: 10.1177/030098587401100108.

Naumann R K, Ondracek J M, Reiter S, Shein-Idelson M, Tosches M A et al. (2015) The reptilian brain. Current Biology, 25 (8), R317–21. DOI: 10.1016/j.cub.2015.02.049.

Ni C, Zhang D, Beyer L A, Halsey K E, Fukui H et al. (2013) Hearing dysfunction in heterozygous Mitf(Mi-wh) /+ mice, a model for Waardenburg syndrome type 2 and Tietz syndrome. Pigment Cell & Melanoma Research, 26 (1), 78–87. DOI: 10.1111/pcmr.12030.

Northcutt R G (2013) Variation in reptilian brains and cognition. Brain, Behavior and Evolution, 82 (1), 45–54. DOI: 10.1159/000351996.

Öfner S, Haugwitz T, Hollandt T, Türbl T, Baur M (2018) Qualzuchten bei Reptilien – Versuch einer Definition. In: 49. Arbeitstagung der AG Amphibien und Reptilienkrankheiten (Arbeitsgemeinschaft der Deutschen Gesellschaft für Herpetologie und Terrarienkunde). Leipzig

Olori J C (2010) Digital Endocasts of the Cranial Cavity and Osseous Labyrinth of the Burrowing Snake Uropeltis woodmasoni (Alethinophidia: Uropeltidae). Copeia, 2010 (1), 14–26. DOI: 10.1643/CH-09-082.

Palci A, Hutchinson M N, Caldwell M W, Lee M S Y (2017) The morphology of the inner ear of squamate reptiles and its bearing on the origin of snakes. Royal Society Open Science, 4 (8), 170685. DOI: 10.1098/rsos.170685.

Pees M (2015) Leitsymptome bei Reptilien. Diagnostischer Leitfaden und Therapie. Enke (kleintier konkret Praxisbuch). Stuttgart.

Pujol R, Rebillard M, Rebillard G (1977) Primary neural disorders in the deaf white cat cochlea. Acta oto-laryngologica, 83 (1-2), 59–64. DOI: 10.3109/00016487709128813.

Résière D, Olive C, Kallel H, Cabié A, Névière R et al. (2018) Oral Microbiota of the Snake Bothrops lanceolatus in Martinique. In: International Journal of Environmental Research and Public Health, 15 (10). DOI: 10.3390/ijerph15102122.

Rieppel O, Kley N J, Maisano J A (2009) Morphology of the skull of the white-nosed blindsnake, Liotyphlops albirostris (Scolecophidia: Anomalepididae). Journal of Morphology, 270 (5), 536–557. DOI: 10.1002/jmor.10703.

Rooney J R (1972) Etiology of the wobbler syndrome. Modern Veterinary Practice, 53 (9), 42.

Rose M P, Williams D L (2014) Neurological dysfunction in a ball python (Python regius) colour morph and implications for welfare. In: Journal of Exotic Pet Medicine, 23 (3), 234–239. DOI: 10.1053/j.jepm.2014.06.002.

Smith S D, Kelley P M, Kenyon J B, Hoover D (2000) Tietz syndrome (hypopigmentation/deafness) caused by mutation of MITF. Journal of Medical Genetics, 37 (6), 446–448. DOI: 10.1136/jmg.37.6.446.

Spaul G, Palmer A, Allsopp J, Hughes-Parry E (1980) Wobbler syndrome (cervical stenosis) in a Percheron colt. Veterinary Record, 107 (15), 362. DOI: 10.1136/vr.107.15.362-a.

Starck J M, Weimer I, Aupperle H, Müller K, Marschang R E et al. (2015) Morphological Pulmonary Diffusion Capacity for Oxygen of Burmese Pythons (Python molurus): A Comparison of Animals in Healthy Condition and with Different Pulmonary Infections. Journal of Comparative Pathology, 153 (4), 333–351. DOI: 10.1016/j.jcpa.2015.07.004.

Starck J M, Schrenk F, Metscher B, Schröder S, Pees M (submitted) Malformations of the scculus and the semicircular canals in spider morph pythons.

Tanchev S, Zhelyazkov E, Philipov J, Semerdjiev V, Paskaslev M et al. (2011) Phenotypic traits of skeletal anomalies observed in inbred rabbits. Revue de Médecine Vétérinaire, 162 (3), 150–153.

Tang W, Zhu G, Shi Q, Yang S, Ma T et al. (2019) Characterizing the microbiota in gastrointestinal tract segments of Rhabdophis subminiatus: Dynamic changes and functional predictions. MicrobiologyOpen, e789. DOI: 10.1002/mbo3.789.

Velo-Antón G, Guilherme Becker C, Cordero-Rivera A (2011) Turtle carapace anomalies. The roles of genetic diversity and environment. PloS One, 6 (4), e18714. DOI: 10.1371/journal.pone.0018714.

Waardenburg P J (1951) A New Syndrome Combining Developmental Anomalies of the Eyelids, Eyebrows and Nose Root with Pigmentary Defects of the Iris and Head Hair and with Congenital Deaf-ness. The American Journal of Human Genetics, 3 (3), 195–253.

Whitfield T T (2015) Development of the inner ear. Current Opinion in Genetics & Development, 32, 112–118. DOI: 10.1016/j.gde.2015.02.006.

Zhang B, Ren J, Yang D, Liu S, Gong X (2019) Comparative analysis and characterization of the gut microbiota of four farmed snakes from southern China. PeerJ, 7, e6658. DOI: 10.7717/peerj.6658.

